# Desiccation-induced DNA damage facilitates drug resistance in *Mycobacterium tuberculosis*

**DOI:** 10.1101/2025.06.04.657859

**Authors:** C. D. Brown, B. M. Lee, H. Liu, A. Wu, A. Tellez, H. Zou, P. Singh, K. Saito, S. Mishra, M. Brown, A. Saleh, N. Odjourian, M. Cristaldo, M. Gan, Q. Liu, M. Gengenbacher, S. Darst, E. Campbell, C. Nathan, K. Y. Rhee

## Abstract

*Mycobacterium tuberculosis* (Mtb) is an obligate human pathogen that depends on its ability to spread from host-to-host to survive as a species. Yet, knowledge of transmission-specific traits remains lacking. Here, we report the discovery of a specific adaptive response to desiccation, a stress intrinsically linked to the generation of the aerosol droplets within which Mtb transmits. We show that desiccation inflicts oxidative damage and activates Mtb’s DNA repair responses but that this repair is imperfect and results in mutations. We further show that activation of these DNA repair responses is accompanied by increased expression of the transcription-coupled repair factor, *mfd,* but that this expression serves to buffer the fitness cost of specific resistance-conferring mutations in *rpoB*, the target of the frontline drug rifampin, rather than to facilitate transcription-coupled DNA repair. Silencing *mfd* during aerosolization impairs survival of strains harboring the rifampin resistance allele S450L. This function is further supported by whole genome sequence data from over 50,000 clinically circulating strains. These studies indicate that Mtb has evolved transmission-specific stress responses that have enabled it to leverage desiccation-induced DNA damage as a potential source of genetic diversification and drug resistance.

## INTRODUCTION

Transmission equips microbes to exert both individual and population level impact (*1, 2*). Control of most infectious diseases thus requires a combination of therapeutic and transmission blocking interventions. Among current infectious diseases, tuberculosis (TB) ranks second only to measles in transmissibility and is the leading cause of deaths due to a single microbial agent (*3*). Unlike most microbial pathogens, *Mycobacterium tuberculosis* (Mtb), the etiologic agent of TB, infects humans as its only known natural host and reservoir (*4–6*). This both makes TB a theoretically eradicable disease and highlights transmission as an obligatory phase of the Mtb life cycle. The essential and repetitive nature of transmission raises the possibility that Mtb may have evolved transmission-specific traits. Knowledge of such traits could serve as the basis for novel drug- and vaccine-based transmission blocking strategies.

Both experimental and epidemiologic data indicate that Mtb is transmitted through air, with its most infectious forms contained in aerosol particles small enough to reach and be retained within the terminal airways and alveoli (*7–11*) (typically between 1-5 microns in aerodynamic diameter) (Fig 1A). The formation of such particles, termed droplet nuclei, is accompanied by a loss of free extracellular water, making desiccation a mechanistically intrinsic feature of TB transmission that is independent of the specific conditions of the external environment (*12–16*). Here, we report the discovery of a specific, adaptive response to desiccation in Mtb. This work not only sheds light on the biology of the most understudied phase of the Mtb life cycle but also lays the groundwork for novel potential transmission blocking strategies.

**Figure 1:**
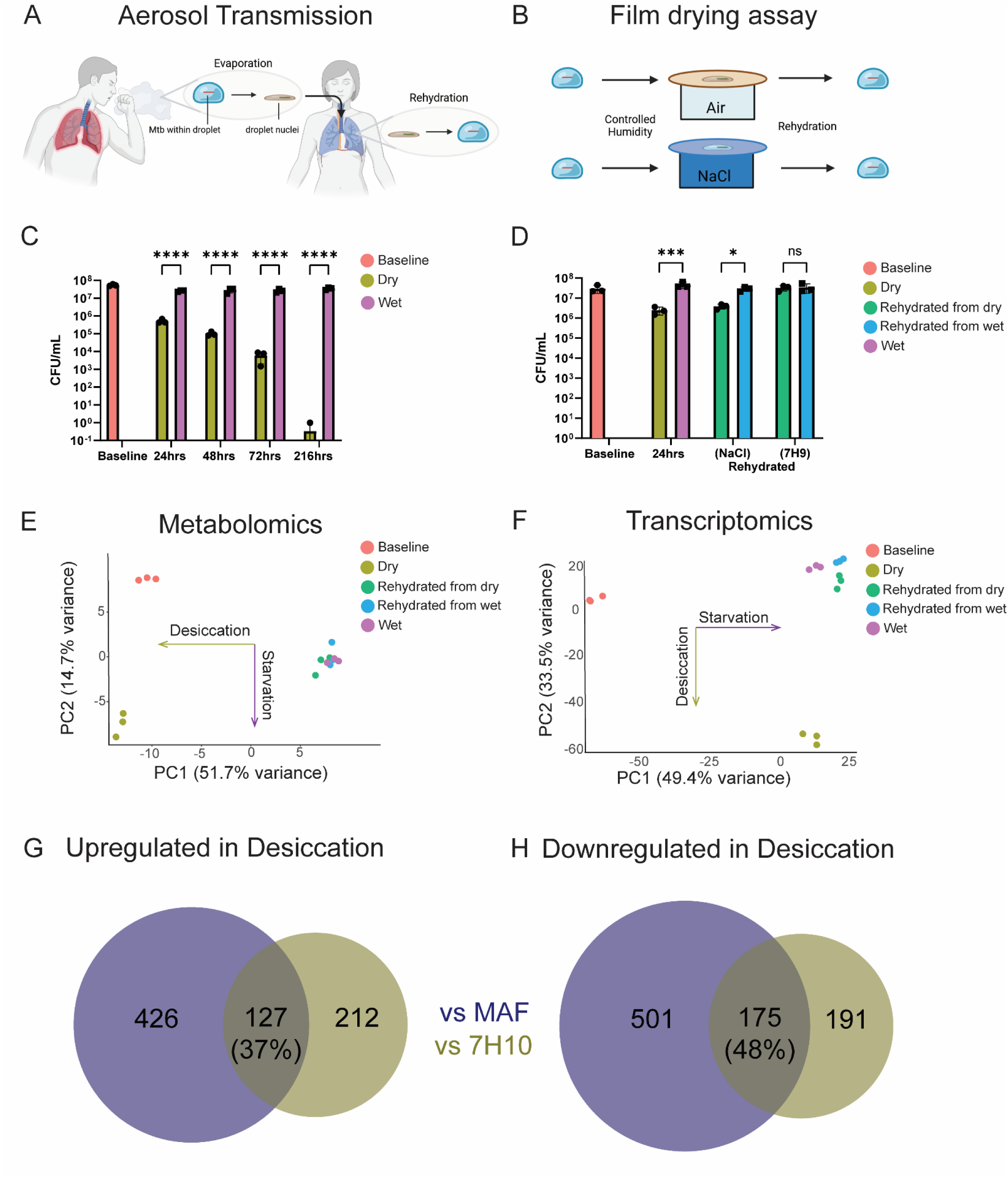
Modeling the desiccation component of aerosol transmission in Mycobacterium tuberculosis. A) An infected host expels Mtb-laden droplets which evaporate to form droplet nuclei; subsequently, these particles are inhaled by a susceptible individual. B) Schematic of drying assay to mimic a desiccation-rehydration cycle. Cells deposited on a sterile filter are allowed to dry out in temperature- and humidity-controlled air and compared against control condition left in contact with a 200 mM NaCl reservoir. C) CFU recovery over 1 week of drying compared to wet (saline) controls. D) CFU recovery following 24 hours of desiccation then rehydration into NaCl or 7H9 media. E) Principal components analysis (PCA) plot of 3 biological replicates from one of two independent experiments comparing baseline (red) metabolome to 24 hours of desiccation (gold), saline control (purple), and rehydration into 200 mM sodium chloride (green and blue). F) PCA plot of 3 biological replicates from one of two independent transcriptomics experiments with batch correction comparing baseline (red) transcriptome to 24 hours of desiccation (gold), saline control (purple), and rehydration into 200 mM sodium chloride (green and blue). G) Venn diagrams prepared using DeepVenn (*55*) of genes with fold change expression >2 (adjusted p-value < 0.05, Benjamini-Hochberg multiple hypothesis correction) or H) Fold change < −2 in Dry vs MAF-adapted (blue) or 7H10-adapted (gold) Mtb. MAF: Model aerosol fluid(*16*). Statistical significance calculated by two-way ANOVA with Tukey’s multiple hypothesis correction, P <.05 (*), .01 (**), .001 (***) or .0001 (****).

## RESULTS

Adapting a previously developed filter culture system that preserves Mtb’s native cell envelope and enables robust experimental modeling of desiccation via evaporative water loss at the air interface (*15, 17, 18*), we characterized Mtb survival following incubation in humidity-controlled air in the presence and absence of access to free water (provided in the form of a contiguous reservoir of saline or air) (Fig 1B). Incubation in air, but not atop saline, revealed a time-dependent multi-log decline in culturability as reported by growth on agar media (Fig 1C) or by enumeration in liquid media by the method of limiting dilution (*19, 20*) (supplemental Fig S1A). This decline could be arrested following incubation in saline, and almost completely reversed with incubation in 7H9 media for 24h prior to plating on agar media (Fig 1D). Recovery yield was not simply due to osmolality of the rehydration liquid (supplemental Fig S1B). Change in culturability, rather than viability, could also be observed across a range of lower humidities (supplemental Fig S1C).

Seeking to further characterize the molecular basis of Mtb’s response to desiccation, we analyzed its transcriptional and metabolic profiles following incubation in ambient air, saline, or desiccation followed by rehydration in saline. Principal components analyses of both metabolomes and transcriptomes independently revealed a striking and unsupervised association between the 2 largest axes of variation and experimental conditions corresponding to nutrient starvation (associated with transfer into saline or air) and desiccation (associated with transfer into and from air), respectively (Fig 1E-F). Focusing first on the transcriptome, we observed a total of 339 and 366 bioinformatically annotated transcripts that exhibited a statistically significant 2-fold increase or decrease (*p*_adj_ < 0.05 after applying Benjamini-Hochberg multiple hypothesis correction) in abundance, respectively, following desiccation (Supplemental Table S1). Consistent with the expected impact of desiccation on intrabacterial osmolality, we observed increased transcript levels of the osmosensory signaling protein Rv0516c, though only limited overlap with the downstream *sigF* regulon it regulates, indicating related but distinct responses to desiccation and osmotic stress (Supplemental Table S1) (*21*). Of the approximately half of desiccation-responsive transcripts bearing bio-informatic annotations in our saline model, we noted a striking enrichment among accumulated transcripts for genes associated with lipid metabolism (59, 16%), toxin-antitoxin pairs (37, 12%), oxidative stress (25, 7%), sulfur metabolism (12, 4%) and DNA repair (10, 3%) and genes associated with translation (41, 13%), purine metabolism (10, 3%) and oxidative phosphorylation (14, 5%) among depleted transcripts (Table 1, Supplemental Table S3). Using a recently developed in vitro model intended to mimic the biochemical and atmospheric conditions encountered by Mtb prior to and during its aerosolization from a necrotic cavity (*16*), we further noted an overlap of 127 (37%) upregulated genes and 175 (48%) downregulated genes (Fig 1G-H, Supplemental Table S2) with similar functional annotations, indicative of a core desiccation transcriptome.

**Table 1:** Significantly upregulated and downregulated genes during desiccation compared to saline-wetted control cells. Genes are considered significantly upregulated if log2 fold change in gene expression was > 1 and adjusted p-value < 0.05 with Benjamini-Hochberg multiple hypothesis correction.

|  | Regulated genes during desiccation (vs saline control) |
| --- | --- |
| **Upregulated** |  |
| Redox | *ald, bfrB, ccdA, cyp138, ethA, fdxC, fixA, fixB, hemL, nadA, nadB, ndh, nrdH, nrdI, pncB1, ribC, rubA, thiX* |
| Sulfur metabolism | *csd, cysA2, cysM, cysO, cysQ, gshA, metH, metK, mmuM, mshA, mshC, mshD* |
| Oxidative damage | *ahpC, ahpD, egtB, egtC, egtD, egtE, oxyS* |
| DNA damage and repair | *lexA, dinG, mfd, mrr, mutT2, polA, radA, recF, ruvA, uvrC* |
| **Downregulated** |  |
| Translation | *def, fusA2, infA, rplB, rplC, rplD, rplE, rplM, rplO, rplP, rplV, rplW, rpmA, rpmC, rpmG2, rpmJ, rpsC, rpsD, rpsE, rpsH, rpsI, rpsJ, rpsK, rpsM, rpsN1, rpsQ, rpsS, rrf, rrl, rrs, ssr, tuf* |
| Nucleotide metabolism | *add, apt, dfrA, glyA2, hpt, mutT1, purD, purN, purU, purH* |
| Oxidative phosphorylation | *atpA, atpB, atpC, atpE, atpF, ctaD, frdA, frdB, frdC, frdD, nuoA, ppa, ppk1, ppk2* |

Studies of other bacteria, yeast and plants have linked desiccation to oxidative stress through a variety of mechanisms. These include a loss of membrane integrity and disruption of the respiratory chain resulting in the accumulation of superoxide ions (O_2_ ^−^), and more widespread alterations in protein conformation and stability that may inactivate proteins involved in anti-oxidant defense (*22–26*). Using a fluorogenic probe of oxidative stress, we observed a time-dependent increase in intracellular reactive oxygen species (ROS) in desiccated, but not saline-treated, Mtb (Fig 2A). Despite this progressive accumulation of reactive radicals, desiccated cells exhibited steady levels of Mtb’s main antioxidant mycothiol by strongly upregulating the biosynthetic genes *mshABCD* (Fig 2B). Nonetheless, the overall increase in ROS was accompanied by extensive oxidative damage as evidenced by a marked rise in 8-oxoguanine (Fig 2C) and lipid peroxides (Fig 2D). Accumulations of oxidized species were specific to desiccation and did not occur in cells held in saline for 24 hours (Fig 2D, supplemental Fig S2). Levels of 8-oxo-2’-deoxyguanosine additionally rose during rehydration, indicative of post-desiccation genomic repair (e.g. glycolytic removal of oxidized bases from the genome and selective hydrolysis of 8-oxo deoxyguanosine triphosphate, dGTP, to deoxyguanosine monophosphate, dGMP) (Fig 2C). Extended desiccation of Mtb revealed a time-dependent increase in double stranded DNA breaks compared to hydrated but nutrient starved controls, as reported by TUNEL staining (Fig 2E).

**Figure 2:**
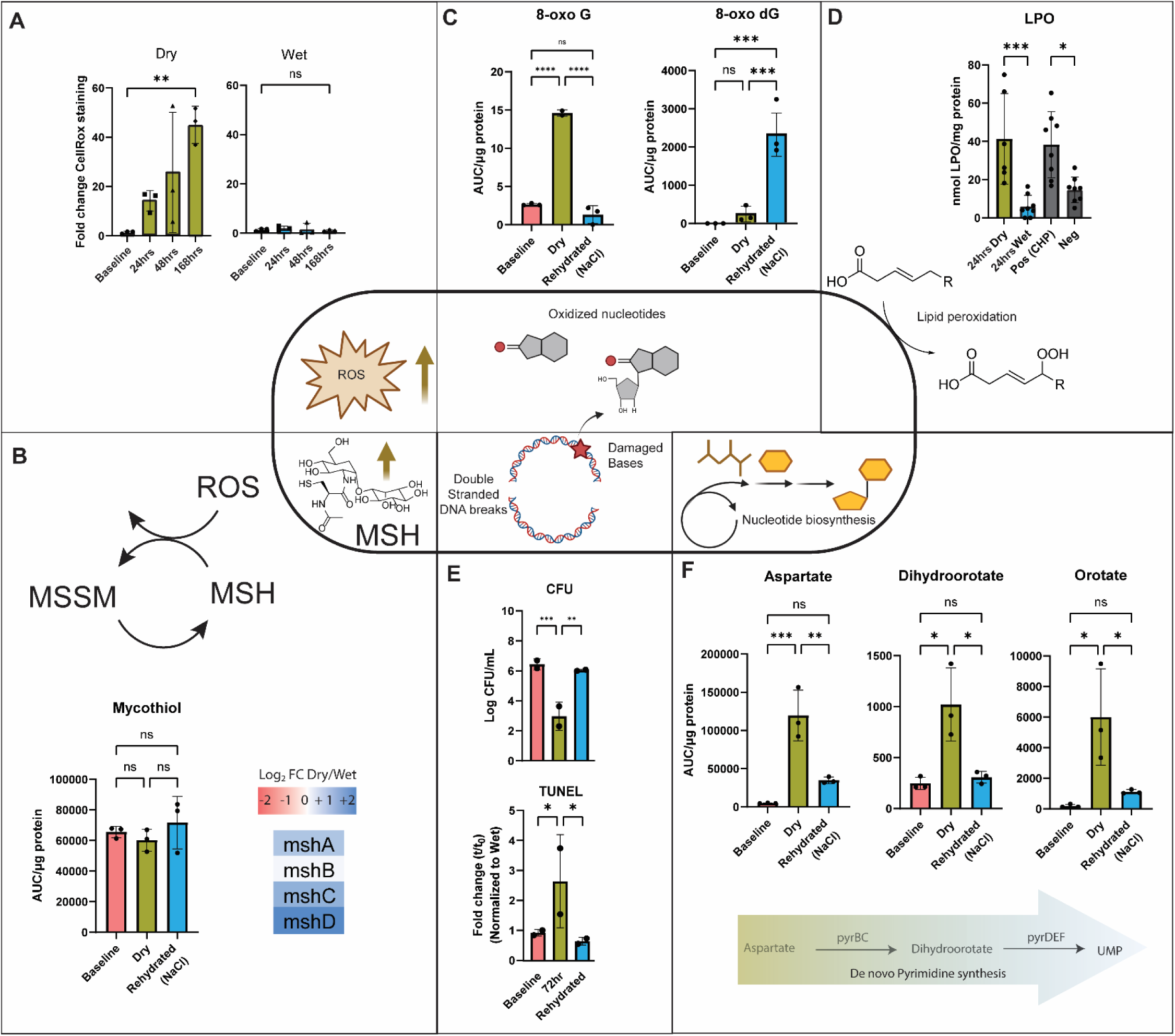
Cells sustain oxidative damage and reversibly adapt to a desiccation-rehydration cycle. A) Cellrox green assay of desiccated versus wet control cells over a 1-week incubation quantified on a BD FACSCelesta Flow cytometer with excitation/emission parameters 488 [515/20] nm normalized to unstained controls. B) Targeted metabolomics of mycothiol levels and RNA Seq gene expression of mycothiol biosynthetic genes (Dry vs Wet). C) Targeted metabolomics of oxidized bases and nucleotides. D) Lipid peroxide abundance quantified by LPO assay of chromogen absorbance at 500 nm normalized to sample protein content by Bradford assay. E) CFU and TUNEL assay of dried and rehydrated cells. Double stranded breaks detected by quantification of FITC-labeling on a BD FACSCelesta Flow cytometer with excitation/emission parameters 488 [515/20] nm normalized to unstained controls. F) Targeted metabolomics of de novo pyrimidine biosynthetic intermediates. All metabolite measurements include three biological replicates from 1 of 2 independent experiments, were quantified by total ion count from an LC-MS assay and normalized to protein content using a Bradford assay. Statistical significance calculated by two-way ANOVA with Tukey’s multiple hypothesis correction, P <.05 (*), .01 (**), .001 (***) or .0001 (****), ns = not significant. MSH: Mycothiol, MSSM: Mycothiol disulfide, TUNEL: terminal deoxynucleotidyl transferase dUTP nick-end labeling, UMP: Uridine-5’-monophosphate

DNA damage notwithstanding, desiccated Mtb exhibited the ability to recover as much as an apparent 3 log_10_ loss of culturability, as reported by colony forming units, after 3 days of desiccation if first allowed to incubate briefly in 7H9 media for 24h prior to enumeration on agar media (Fig 2E). This CFU recovery correlated with a concordant decrease in TUNEL staining indicative of rehydration associated repair (Fig 2E). Consistent with the need for *de novo* nucleotide biosynthesis (*27*), metabolic profiling revealed desiccation-induced accumulations of aspartate, dihydroorotate and orotate, metabolic intermediates of *de novo* pyrimidine biosynthesis, that were reversed upon rehydration and linked to an increase in levels of the pathway end product, uridine 5’-monophosphate (UMP) (Fig 2F, supplemental Fig S2). These increases were not observed in non-desiccated, nutrient starved controls incubated in saline (Supplemental Fig S2).

Closer analysis of the Mtb transcriptome following 24h of desiccation similarly revealed increased transcript levels of genes of the nucleotide excision repair *(uvrA/Rv1638, uvrC/Rv1420, polA/Rv1629*), transcription coupled repair (*mfd/Rv1020*), and base excision repair (*mutT2/Rv1160, nth/Rv3674c*) pathways, genes involved in DNA strand break detection and repair (*recF/Rv0003, lexA/Rv2720, radA/Rv3585*, *ruvA/Rv2593c, xseA/Rv1108c*) (Table 1, Fig 3A, 3B). Rehydration triggered further increases in expression of additional DNA repair systems including components of the base excision repair system that specifically recognize mutagenic oxidized purines (*mutM/fpg*/*Rv2924c*) DNA ligases (*ligB/Rv3062*) and the error prone DNA polymerase *dinB1*/*Rv1537* (Fig 3C).

**Figure 3:**
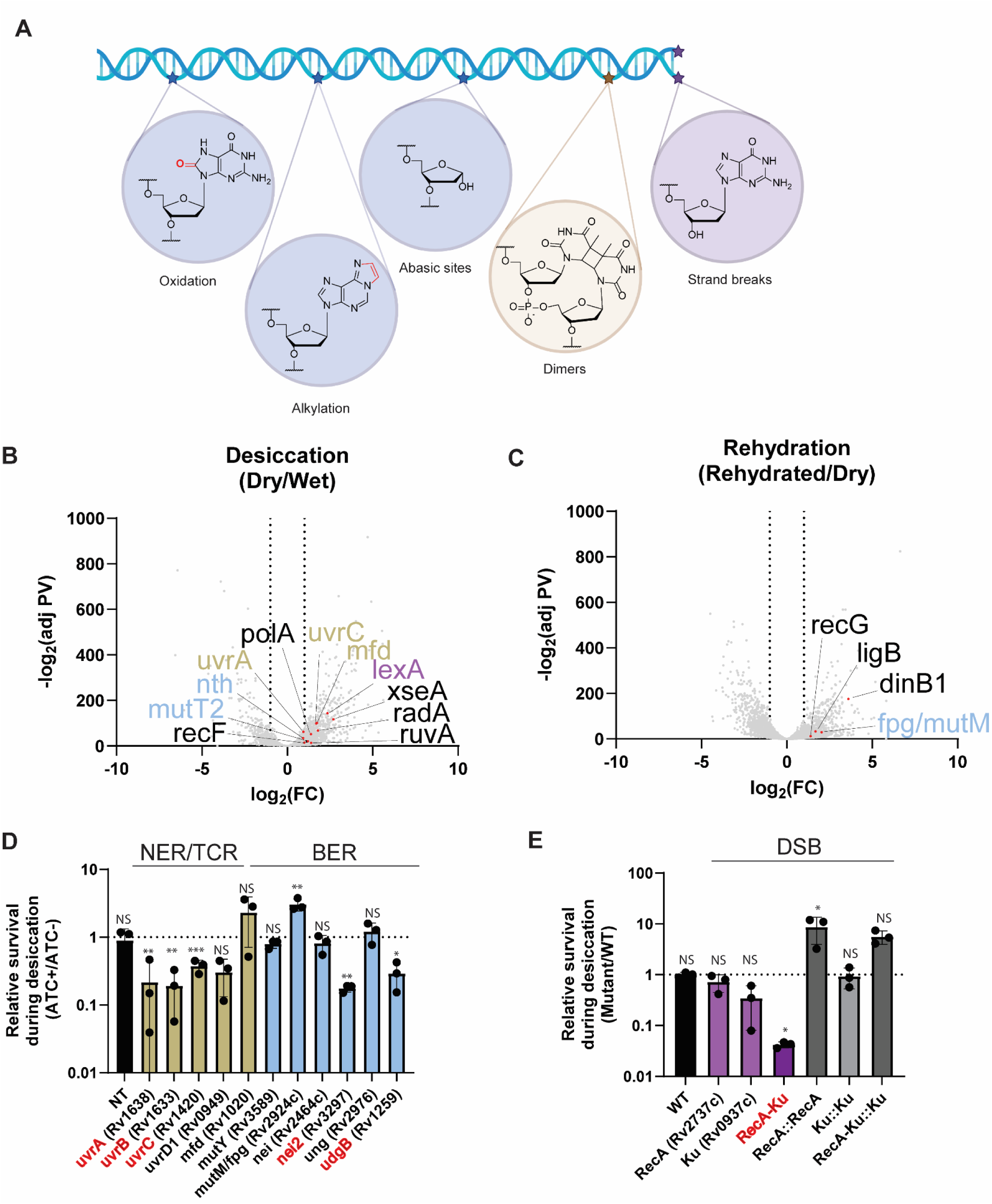
A multi-pronged DNA damage and repair transcriptional response is critical during desiccation-rehydration. A) Examples of chemically modified DNA bases color coded by repair processes that recognize them. Base excision repair (BER), blue; nucleotide excision repair (NER) or transcription coupled repair (TCR), gold; double stranded breaks (DSB), purple. B) RNA Seq volcano plot of annotated transcripts comparing cells desiccated for 24 hours to wet (saline) controls. Genes involved in NER/TCR, BER and DSB repair in gold, blue and purple, respectively. Additional polymerases, helicases and ligases in black. C) RNA Seq volcano plot of annotated transcripts comparing cells desiccated for 24 hours to cells rehydrated (into saline). D) Screen of CRISPRi knockdown strains after 24 hours of desiccation targeting NER, TCR and BER pathways plotted as a ratio of the relative survival of the knockdown compared to the ATC-control. E) Screen of DSB repair mutant strains after 24 hours of desiccation plotted as a ratio of the relative survival of the strain compared to wild type. See methods for details. Statistical significance calculated by 2-tailed, unpaired student’s t-test between ATC+/ATC-(D) or mutant/WT (E), P <.05 (*), .01 (**), .001 (***) or .0001 (****), ns = not significant. Strains with a significant decrease in viability highlighted in red. NT = non-targeting CRISPRi strain, ATC: anhydrotetracycline, WT: Wild type (Erdman).

Seeking to define the specific DNA repair mechanisms required to survive desiccation-induced DNA damage, we designed and tested a library of CRISPRi knockdown strains targeting elements of nucleotide excision repair (NER), transcription coupled repair (TCR) and base excision repair (BER) pathways, as well as key genes involved in DNA strand break repair (*28, 29*). Scoring on the relative survival of each strain to itself when silenced (or not) after 24 hours of desiccation, we recovered fewer CFU with silencing of specific components of the NER and BER pathways, potentially indicative of non-redundant roles associated with repair of specific types of DNA damage (Fig 3D, supplemental Fig S3). In contrast, we observed a survival defect only in a *recA*-*ku* double knockout, indicative of their functional redundancy in repair of otherwise lethal double stranded DNA breaks (Fig 3E). These findings thus collectively provide evidence of an integrated adaptive response to desiccation-induced DNA damage.

Given Mtb’s capacity to withstand and recover from desiccation-induced DNA damage, we also probed for evidence of desiccation-induced mutagenesis. Using rifampin resistance as a phenotypic readout of point mutations, we observed a 50-100 fold increase in the frequency of rifampin resistance following desiccation compared to incubation in saline (Fig 4A). This increase was elevated even further upon inhibition of the nucleotide excision repair helicase, *uvrC*, by CRISPRi (supplemental Fig S4A). However, these increases could be extinguished altogether if desiccated Mtb was first allowed to briefly recover in 7H9 for 24 hours prior to plating on agar media (Fig 4A).

**Figure 4:**
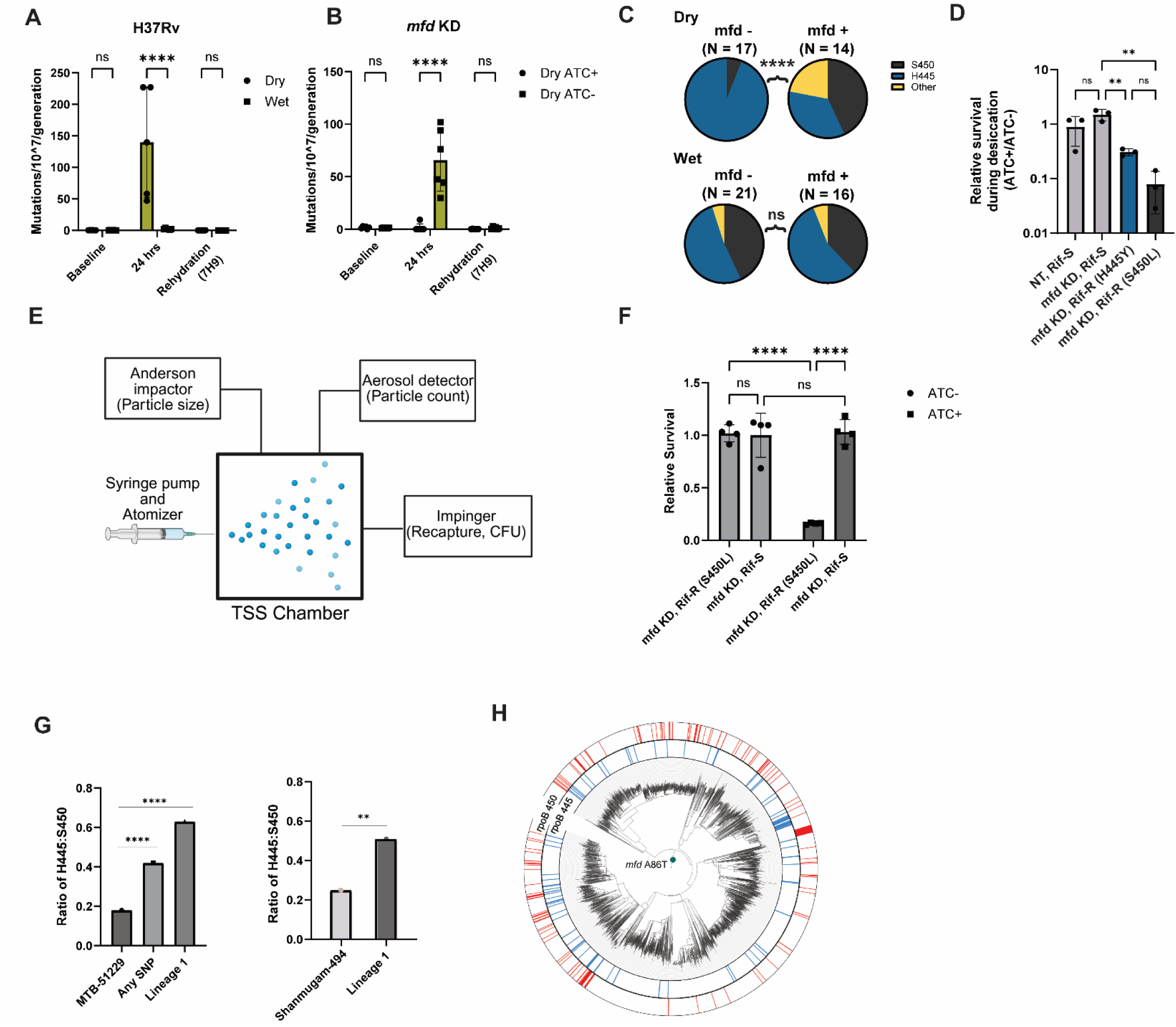
Desiccation-induced mutagenesis promotes rifampin resistance via the activity of Mfd. Rif-R mutation frequency of desiccated cells of laboratory strain H37Rv (A) or mfd knockdown (B) before desiccation, immediately after 24 hours of desiccation, and following 24 hours rehydration after desiccation into nutrient rich media (7H9). Each panel displays 6 biological replicates from 1 of 2 independent experiments. Mutation frequency was calculated using the online fluctuation analysis calculator FALCOR (genomics.brocku.ca/FALCOR) using the Lea-Coulson method. Statistical significance was determined by two-way ANOVA with Tukey’s multiple comparison’s test. C) Rif-R allele frequency from two independent experiments following 24 hours of desiccation +/-inhibition of *mfd*. Statistical significance determined by chi squared analysis. D) Representative experiment from 2 independent assays each with 3 biological replicates of relative survival of mfd-silenced Rif-R strains after desiccation. Statistical significance determined by two-tailed unpaired student’s t-test. E) Schematic of TSS system (Nuritdinov 2025, *submitted*) F) Relative survival of Rif-S (*rpoB* WT) and Rif-R (*rpoB* S450L) *mfd* KD strains after aerosolization in the TSS, ATC: anhydrotetracycline, RIF: rifampin. G) Ratio of H445:S450 Rif-R mutants in MTB-51229 global lineage data set (*41*), subset analysis, and from Shanmugam et. al (*42*). Statistical significance determined by Fisher’s exact test. H) Phylogenic tree of Lineage 1 strains and I) clade L4.1.1 within MTB-51229. rpoB H445 SNP (blue) and rpoB S450 SNP (red). P <.05 (*), .01 (**), .001 (***) or .0001 (****), ns = not significant. ATC: Anhydrotetracycline, TSS: Transmission simulation system

The foregoing “mutation frequency decline” focused our attention on the transcription-coupled DNA repair factor Mfd, transcript levels of which increased with desiccation and decreased with rehydration into saline (Supplemental Tables S1, S3, Fig 3B). Mfd is a highly conserved DNA translocase that was first identified based on its phenotypic ability to regulate the reversion of auxotrophic mutants of *E. coli* following ultraviolet irradiation (*30, 31*). Mfd has since been shown to facilitate nucleotide excision repair at sites of DNA damage by displacing stalled RNA polymerase complexes (*32–34*). Recent work has expanded the physiologic range of Mfd’s activities to include more general housekeeping roles in RNA polymerase processivity, and in some cases somewhat paradoxically, inhibition of DNA repair (*35, 36*). Ragheb et al. additionally reported a specific role for Mfd in promoting antibiotic resistance conferring mutations(*37*). We found that CRISPRi-mediated transcriptional silencing of *mfd* had no impact on survival of the aggregate population of Mtb prior to or during desiccation (supplemental Fig S4B). Scoring on rifampin resistance however revealed that silencing of *mfd* suppressed the frequency of rifampin resistance following desiccation (Fig 4B), suggesting that Mfd either antagonized repair of desiccation-induced DNA lesions or promoted the fitness of rifampin resistant (Rif-R) Mtb. Scoring on ciprofloxacin resistance, which similarly arises from point mutations in its target (DNA gyrase), we observed a similar increase in the frequency of mutants following desiccation (supplemental Fig S4C). Unlike the case for rifampin however, this increase was unaffected by silencing of *mfd*, supporting a role for *mfd* in the fitness of rifampin resistant mutants (supplemental Fig S4C).

Targeted sequencing of the 81 bp rifampin resistance determining region (RRDR) (that mediates 96% of all clinically observed rifampin resistance) from desiccation-induced rifampin resistant strains revealed a striking and near complete replacement of strains harboring the clinically most prevalent rifampin resistance allele, S450L, by strains harboring the second most prevalent allele, H445X, in *mfd*-silenced strains (*3, 38*) (Fig 4C, Supplemental Table S4). Gagneux *et al*. reported that Rif-R strains harboring either drug resistance allele exhibited fitness defects when competed against drug sensitive strains in vitro (*39*). Eckartt *et al*. more recently showed that the observed fitness defect of drug-resistant strains is the product of both the ‘direct’ fitness cost of the isolated drug resistance allele and the activity of factors capable of compensating for specific physiologic defects it incurs (*40*). We tested the impact of *mfd* silencing on survival of S450L Rif-R Mtb prior to, during and following desiccation, and observed a selective 1 log_10_ CFU defect in S450L, but not H445Y, Rif-R Mtb strains following desiccation and rehydration (supplemental Fig S4D-E). Plotted as a function of relative survival, drug-sensitive *mfd* knockdown strains had no survival defect, whereas Rif-R strains exhibited an 8- and 28-fold decline in viability for H445Y and S450L, respectively (Fig 4D).

To further validate the relevance of our findings, we made use of a newly developed precision aerosol generation system, termed the transmission simulation system (TSS) (Fig 4E) (Nuritdinov 2025, *under review*). The system allows for controlled generation of aerosol particles of defined size and in-flight monitoring of particle size and concentration as well as particle recapture by coupling to a liquid impinger (Fig 4E). Using this system, we observed a similar desiccation-specific fitness defect of Rif-R (*rpoB* S450L), but not Rif-S, strains only when *mfd* had been pre-depleted (Fig 4F). These results thus demonstrate a role for *mfd* in promoting the *in vitro* fitness of Rif-R Mtb arising from desiccation-induced mutagenesis.

Seeking clinical evidence of such a compensatory function, we analyzed a previously curated (*41*) database of 51,229 Mtb whole genome sequences containing 10,230 S450L and 1,890 H445Y Rif-R isolates (rpoB445/rpoB450 = 0.18) (Fig 4G). This database also contained 7,062 isolates encoding nonsynonymous mutations in *mfd*. Among *mfd* mutants, 587 harbored the S450L mutation, while 244 encoded the H445Y allele (rpoB445/rpoB450 = 0.42) resulting in a statistically significant enrichment of H445Y mutants compared to that of all Rif-R isolates (p < 0.0001 by Fisher’s exact test) (Fig 4G). We further found that strains belonging to lineage 1 (L1) encoded an ancestral mutation in *mfd* (A86T), which is a lineage-defining mutation. Among 4,766 L1 strains, 138 harbored the H445Y allele, while 220 encoded the S450L rifampin resistance allele (rpoB445/rpoB450 = 0.63) resulting in an even greater, statistical enrichment of H445Y mutants (p < 0.0001 by Fisher’s exact test) (Fig 4G). Phylogenetic reconstruction of H445Y mutational events indicates that the increased prevalence of this mutation in L1 strains is not associated with specific outbreak events; instead, it appears to have accumulated independently in parallel across multiple phylogenetic clades (Fig 4H), suggestive of a specific genetic interaction with *mfd*. Work by Shanmugam et. al. reported a similar trend among 494 circulating strains in India representing global lineages L1, L2, L3 and L4 (*42*). Among the 494 strains analyzed there were 366 Rif-R strains containing 244 S450L and 61 H445Y (rpoB445/rpoB450 = 0.25) (Fig 4G). Within the 211 L1 strains 134 were Rif-R including 72 S450L and 35 H445Y (rpoB445/rpoB450 = 0.51) which was statistically significant (p <0.01 by Fisher’s exact test) (Fig 4G).

Modeling the Mtb Mfd (Mfd_Mtb_) WT and A86T variant using AlphaFold and a previously published cryo-EM structure of E. coli Mfd (Mfd_Eco_) (*43*), we mapped the A86 residue to the uvrB homology domain, corresponding to residue M66 in E. coli (Supplemental Fig S5A, B), and discovered a potential steric impact of a threonine substitution with neighboring residues suggestive of a conformational change (supplemental Fig S5C-D). Awaiting a more complete biochemical and structural characterization of this variant, these findings support a potentially deleterious impact of the A86T mutation on Mfd’s fitness buffering activity on slower RNA polymerase variants such as S450L.

## DISCUSSION

Unlike the case for other microbes, transmission is a feature of both the pathogenicity and physiology of Mtb. Despite its erratic and often fleeting nature, transmission is an obligatory phase of a lifecycle whose repetitive nature may be sufficient to impose a selective pressure on Mtb as a species. Our studies implicate desiccation stress as one such potential pressure. This is because the most transmissible forms of Mtb correspond to aerosol particles with limited free extracellular water content (*9–11*) yet derive from the humid environment of the lung and airways (relative humidity of 30-70%) (*44*) making desiccation an intrinsic, if not invariant, feature of all transmission events. We specifically show that: (i) desiccation incurs a combination of widespread oxidative damage and osmotic stress; (ii) this stress renders Mtb into a phenotypic state that is unable to replicate on agar media but can be almost fully restored if allowed to briefly recover in nutrient-rich media; and (iii) entry and exit from this state is linked to a molecular program that is specifically induced by desiccation but can be reversed with rehydration into saline alone. Transcriptional profiling more specifically revealed a striking upregulation of genes annotated to be associated with oxidative and osmotic stress, lipid remodeling, and tRNA and rRNA cleavage; and linked to downregulation of genes annotated to encode components of the ribosome and tRNA synthetases, and oxidative phosphorylation (Table 1, Supplemental Tables S1-S3). These changes thus demonstrate that desiccation elicits a specific molecular program that consists of an oxidative stress response, active reprogramming/arrest of translation, and inhibition of aerobic respiration.

Our studies further show that desiccation, when not lethal, is mutagenic. Mutagenic diversification is a population-level strategy to ensure success of the species. However, factors that promote genetic diversity must be balanced against those that promote repair. In *B. subtilis,* for example, amino acid auxotroph revertants arising from a stationary phase mutagenesis model require an interplay between *mfd*, *mutM/fpg*, *mutY,* the NER system and error-prone polymerases (*45, 46*). Mtb’s repair response provides further support of a repetitive history of exposure and adaptation to desiccation-induced DNA damage. Our studies indicate that in Mtb mutagenicity is tempered by the NER system component, *uvrC*, and further reflected in the joint essentiality of both single and double stranded DNA repair mechanisms for optimal survival of Mtb under desiccation stress (*29, 47–51*) (Fig 3E). Depending on the extent of opportunity for repair desiccation may thus facilitate a limited degree of genetic diversification among the surviving population of Mtb.

The transcriptional upregulation and role of *mfd* in Mtb’s response to desiccation merits special discussion. This is because the gene-selective, and *uvrC*-independent, impact of *mfd* on the frequency and spectrum of mutations in *rpoB* indicates that *mfd* may facilitate adaptation to, rather than repair of, desiccation-induced DNA damage. This interpretation is supported by the following observations. Rif-R mutations in *rpoB* impair Mtb fitness even in the absence of drug pressure (*39*). Rif-R mutations in *rpoB* have pleiotropic effects on the enzymatic activity of Mtb RNA polymerase (RNAP) (*40*). The most common clinical mutation in *rpoB* (S450L) causes Mtb RNAP to elongate more slowly and terminate more frequently at sites of transcriptional pausing than wild type RNAP, while the H445Y mutation is associated with faster rates of elongation (*40*). It is thus plausible that, in the setting of a damaged genome, the DNA translocase activity of Mfd may specifically help the S450L mutant compensate for even higher rates of pausing and termination caused by its slowed elongation rate. Such a hypothesis is supported experimentally by the desiccation-specific essentiality of *mfd* in an S450L mutant observed in both the filter-mounted drying (Fig 4D) and aerosol survival assays utilizing the transmission simulation system (Fig 4F), and the reduced frequency of S450L mutations observed in L1 strains of Mtb bearing an ancestral mutation in *mfd* (*42*) (Fig 4G-H). Whether or how the ancestral A86T mutation of *mfd* among L1 strains alters the fitness conferring activity of *mfd* is not known. Awaiting future studies on the specific mechanisms by which *mfd* may buffer the fitness cost of other *rpoB* mutations, the desiccation-specific nature and essentiality of *mfd* in the frequency and spectrum of *rpoB* mutations support its role as an adaptive post-mutational trait that has evolved in response to the mutagenic impact of desiccation. Indeed, lineage-specific fitness costs of specific Rif-R conferring *rpoB* mutations have previously been described in which transmission rates of S450L outpace other mutants in L2, but not L4 (*52*). In communities where L1 strains co-circulate with other lineages, the frequency of Rif-R acquisition in L1 lags the other lineages (*42, 53*). Furthermore, recent work indicated positive selection on *mfd* among locally adapting strains on the Tibetan plateau (*54*).

The specific degree of desiccation and DNA damage experienced by Mtb during transmission remain to be defined. However, the inherently desiccating nature of aerosol generation and Mtb’s integrated physiologic response to rehydration provide strong evidence of an evolutionary impact on Mtb physiology. Uncertainties notwithstanding, this work provides evidence of a specific and previously undescribed feature of Mtb physiology that may contribute to the arguably least studied phase of its life cycle and key determinant of its population level impact.

## Acknowledgments

The authors would like to acknowledge Weill Cornell Genomics Core Facility for assistance with RNA Seq bioinformatics, The Flow Cytometry Core Facility for access to cytometers, and Michael Glickman (Memorial Sloan Kettering) for providing the RecA and Ku mutants. Figures were made in GraphPad Prism, Adobe Illustrator, DeepVenn and Biorender.

## Funding

National Institutes of Health grant K08AI148584 (CB)

National Institutes of Health grant P01AI159402 (KR, CN, MG)

National Institutes of Health grant T32AI07613 (CB)

Weill Department of Medicine Fund for the Future (CB)

## Author contributions

Conceptualization: CB, KR

Formal Analysis: CB, KR

Funding Acquisition: CB, KR

Methodology: CB, BL, HL, AW, AT, HZ, KS, SM, KR

Investigation: CB, BL, HL, AW, AT, HZ, KS, SM, PS, MB, AS, FN, JW, MGa, QL, SD, EC

Visualization: CB

Project administration: CB, KR

Resources: KR, MGe, QL

Supervision: KR, CN

Writing – original draft: CB, KR

Writing – review & editing: CB, KR

**Supplementary Materials**

Materials and Methods

Figs. S1 to S5 Tables S1 to S4

---

**Supplemental Figure S1:**
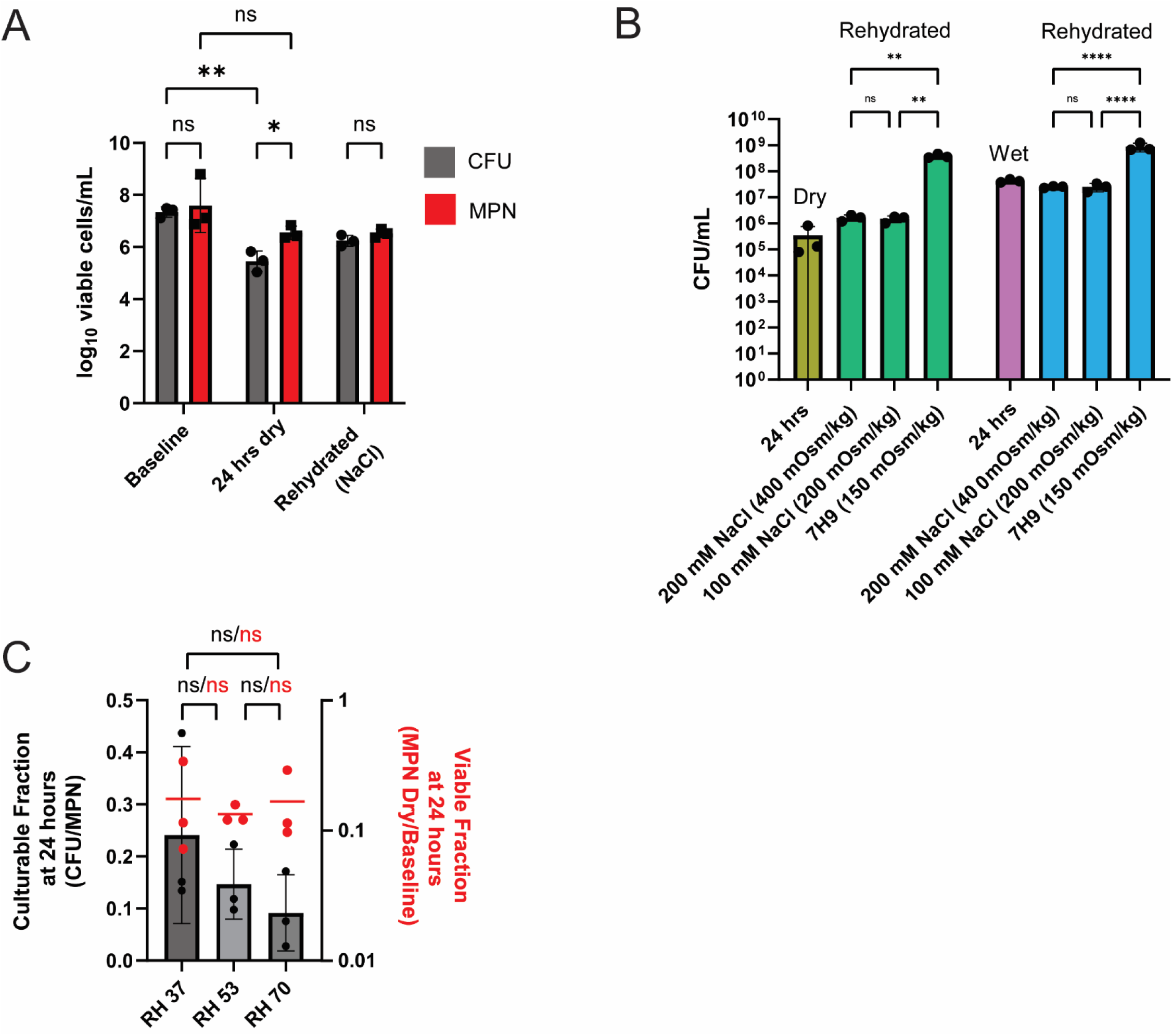
Cell viability under varying osmolality and humidity. A) Enumeration of viable cells by CFU (grey) or MPN (red) after 24 hours at 70% relative humidity (RH). CFU recovery during rehydration after 24 hours of desiccation. Statistical significance by two-way ANOVA with Tukey’s multiple comparison. Culturable fraction by CFU/MPN ratio (grey) and surviving fraction by MPN normalized to baseline biomass (red) across 3 humidities. P <.05 (*), .01 (**), .001 (***) or .0001 (****), ns = not significant. CFU: colony forming units, MPN: most probable number.

**Supplemental Figure S2:**
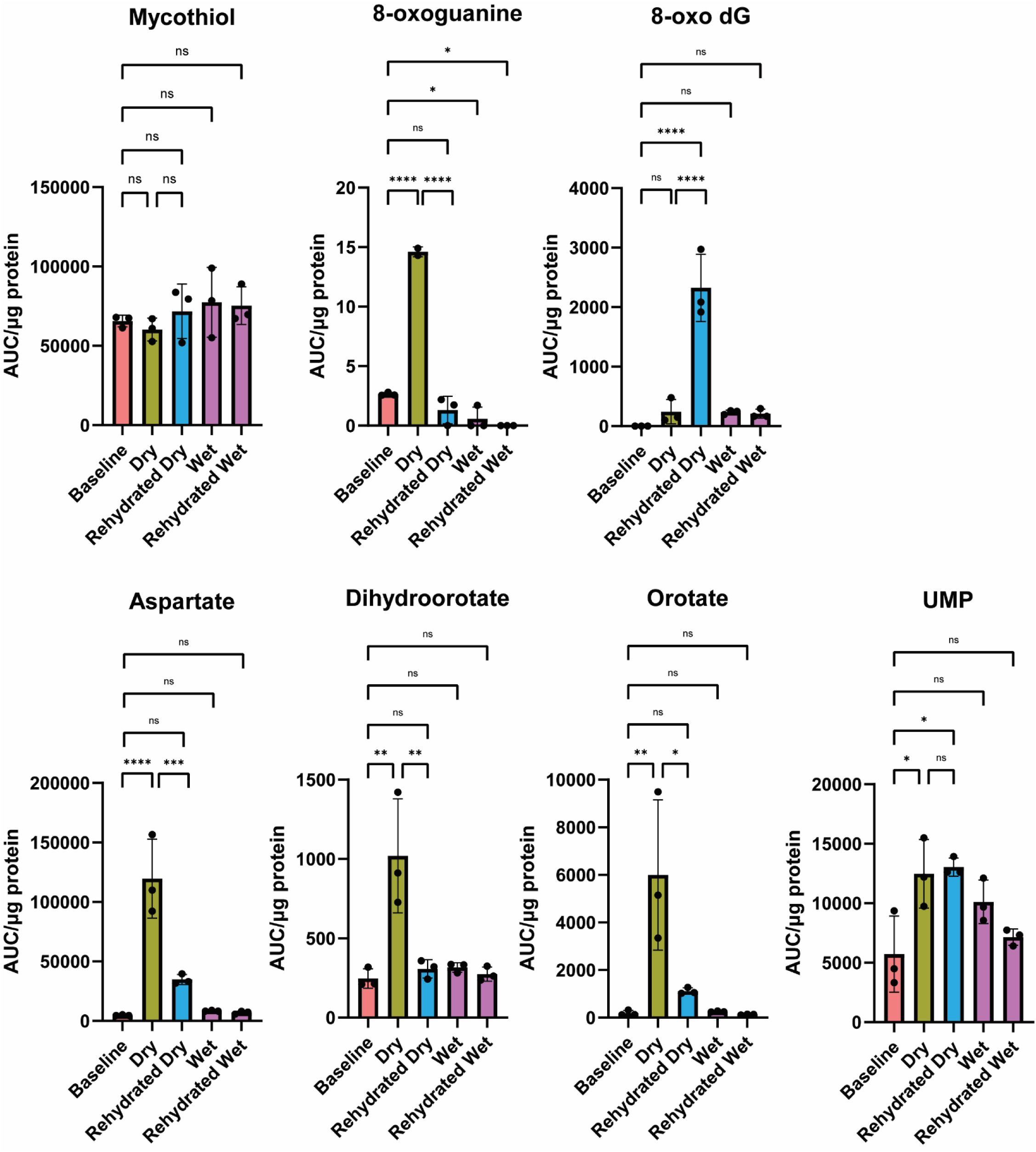
Targeted metabolomics of mycothiol, oxidized nucleotides and de novo nucleotide biosynthetic intermediates quantified by total ion count from LC-MS assay and normalized to protein content by Bradford assay. Biological triplicates from 1 of 2 independent experiments. Statistical significance calculated by one-way ANOVA with multiple hypothesis correction, P <.05 (*), .01 (**), .001 (***) or .0001 (****), ns = not significant.

**Supplemental Figure S3:**
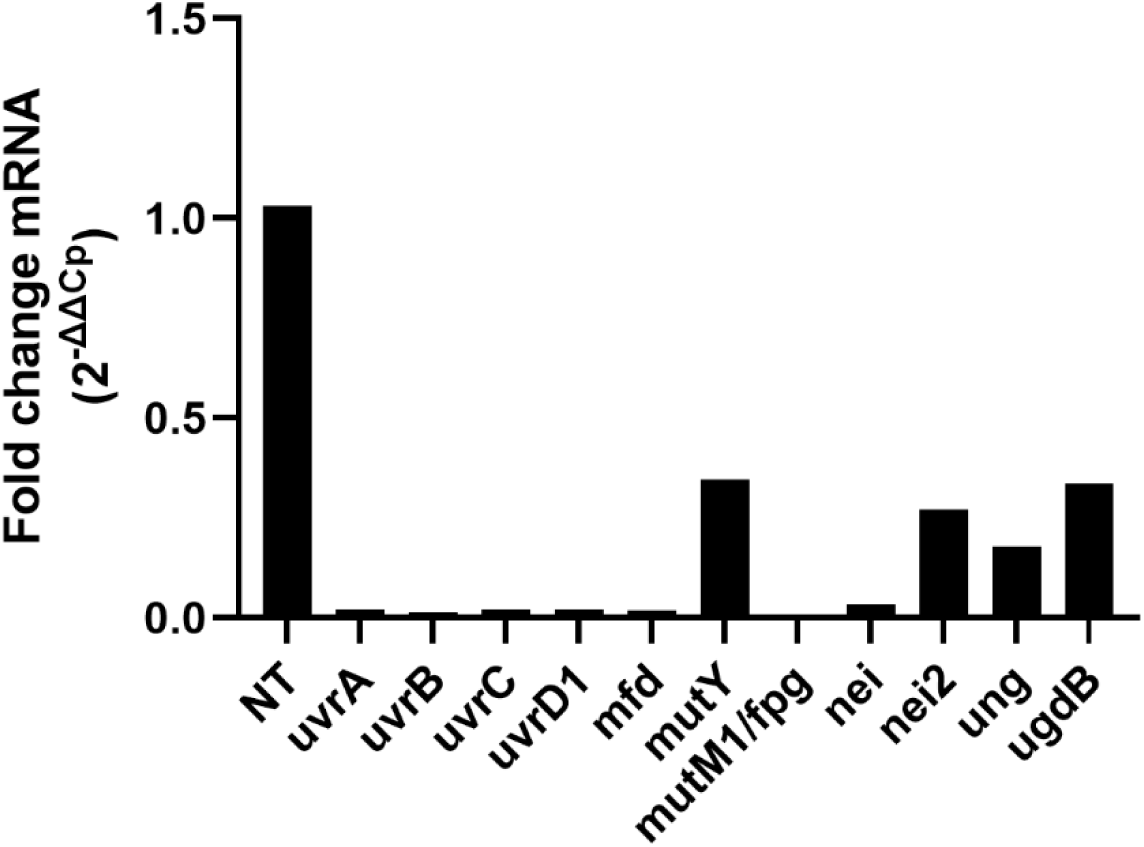
Fold change in gene expression for CRISPRi strains presented in Figure 3. Calculations of ΔΔCp by RT qPCR normalizing the target gene to housekeeping gene *sigA*. NT fold change was calculated for *mfd* expression.

**Supplemental Figure S4:**
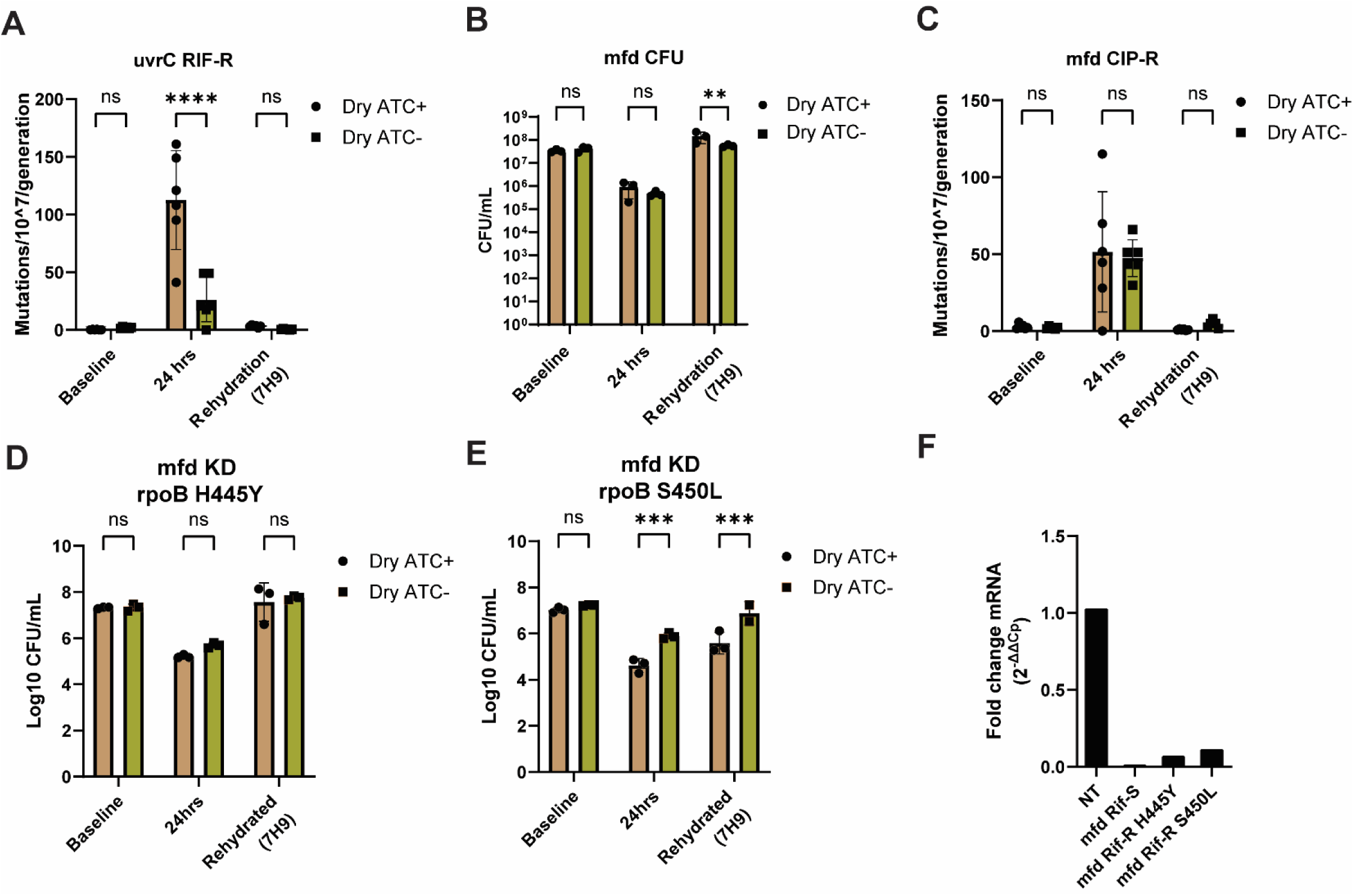
Mutation rate and survival during desiccation for uvrC and mfd CIRSPRi KD strains. A) Mutation frequency for *uvrC* KD during pre-desiccation baseline, 24 hours of drying, and rehydration into nutrient-rich 7H9 media. Mutation frequency was calculated using the online fluctuation analysis calculator FALCOR (genomics.brocku.ca/FALCOR) using the Lea-Coulson method. Statistical significance was determined by two-way ANOVA with Tukey’s multiple comparison’s test. B) Survival in CFU/mL for *mfd* CRISPRi KD during pre-desiccation baseline, 24 hours of drying, and rehydration into nutrient-rich 7H9 media. C) Mutation frequency for *mfd* KD during pre-desiccation baseline, 24 hours of drying, and rehydration into nutrient-rich 7H9 media. Survival in log10 CFU/mL for mfd KD strains harboring rpoB Rif-R mutant alleles H445Y (D) or S450L (E) during pre-desiccation baseline, 24 hours of drying, and rehydration into nutrient-rich media. F) Fold change in mRNA expression upon ATC-induced CRISPRi induction normalized to housekeeping gene *sigA*. Statistical significance by two-way ANOVA with Tukey’s multiple comparison. P <.05 (*), .01 (**), .001 (***) or .0001 (****), ns = not significant. CFU: colony forming units, ATC: Anhydrotetracycline, KD: knockdown, RIF-R: Rifampin resistance, CIP-R: Ciprofloxacin resistance.

**Supplemental Figure S5:**
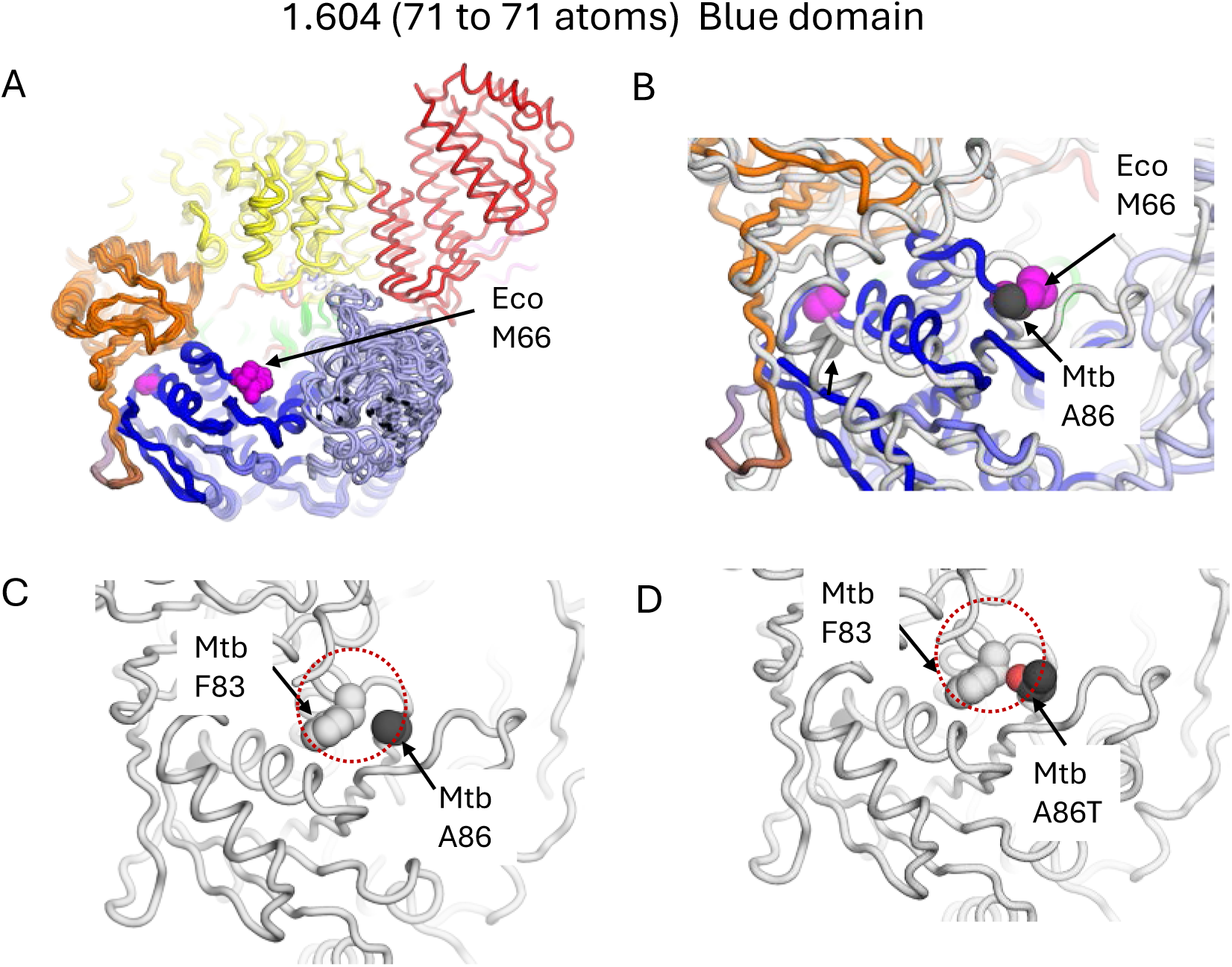
Comparison of the structures of *E. coli* Mfd (Mfd_Eco_) to *M. tuberculosis* Mfd (Mfd_Mtb_). A) Cryo-EM structure of Mfd_Eco_ in L0 (repressed) state centered on domain 1 (D1, Blue) as originally described in Kang et. al.(*43*). Residue Eco M66 (Magenta spheres) corresponds to Mtb A86 B) Alphafold prediction (white ribbon) of Mfd_Mtb_ overlayed on Mfd_Eco_. No local conformational change around Mtb A86 (black spheres) relative to Eco M66 (magenta spheres). The fit across all 71 D1 atoms showed variability with RMSD of 16.0 Å. C) The residue Mtb F83 (grey spheres) in close proximity to Mtb A86. D) Potential steric interactions (red dashed circles) between Mtb F83 and Mtb A86T mutation.

## Supplemental Materials

### Oligonucleotides used in this study

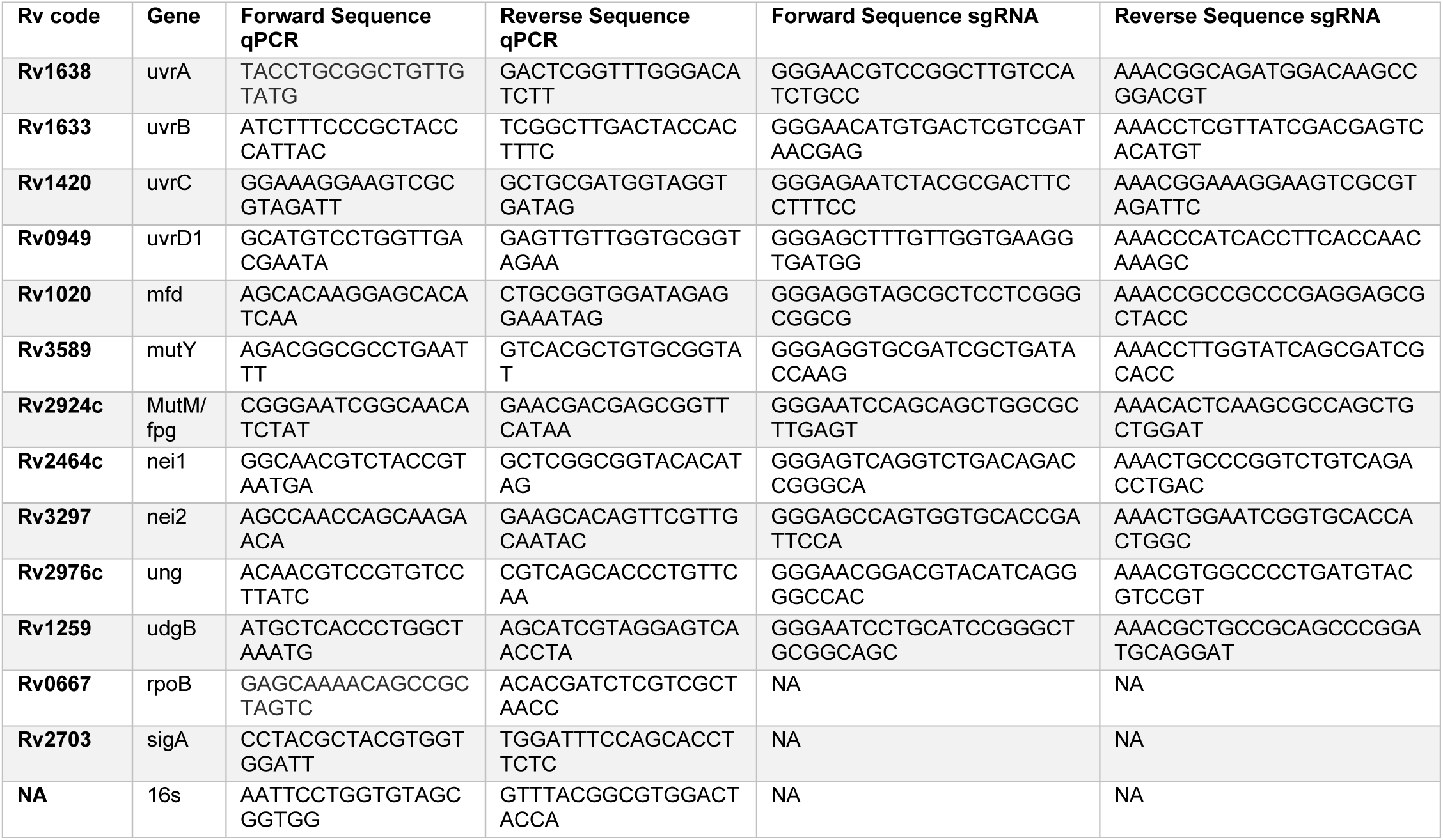

### Strains used in this study

| Strain | Rv number | Description | Source |
| --- | --- | --- | --- |
| **H37Rv** | NA | Laboratory Strain | ATCC |
| **Erdman** | NA | Laboratory strain | ATCC |
| **NT** | NA | Non-targeting CRISPRi strain in H37Rv background | Rock et. al. 2017 |
| **uvrA** | Rv1638 | CRISPRi targeting uvrA in H37Rv background | This work |
| **uvrB** | Rv1633 | CRISPRi targeting uvrB in H37Rv background | This work |
| **uvrC** | Rv1420 | CRISPRi targeting uvrC in H37Rv background | This work |
| **uvrD1** | Rv0949 | CRISPRi targeting uvrD1 in H37Rv background | This work |
| **mfd** | Rv1020 | CRISPRi targeting mfd in H37Rv background | This work |
| **mfd rpoB H445Y** | Rv1020 | CRISPRi targeting mfd in H37Rv background with rpoB H445Y allele | This work |
| **mfd rpoB S450L** | Rv1020 | CRISPRi targeting mfd in H37Rv background with rpoB S450L allele | This work |
| **mutY** | Rv3589 | CRISPRi targeting mutY in H37Rv background | This work |
| **mutM/fpg** | Rv2924c | CRISPRi targeting mutM/fpg in H37Rv background | This work |
| **nei** | Rv2464c | CRISPRi targeting nei in H37Rv background | This work |
| **nei2** | Rv3297 | CRISPRi targeting nei2 in H37Rv background | This work |
| **ung** | Rv2976c | CRISPRi targeting ung in H37Rv background | This work |
| **udgB** | Rv1259 | CRISPRi targeting udgB in H37Rv background | This work |
| **RecA** | Rv2737c | RecA knockout in Erdman background | Heaton et. al. 2014 |
| **Ku** | Rv0937c | Ku knockout in Erdman background | Heaton et. al. 2014 |
| **RecA-Ku** | Rv2737c, Rv0937c | RecA-Ku double knockout in Erdman background | Heaton et. al. 2014 |
| **RecA::RecA** | Rv2737c | Complement | Heaton et. al. 2014 |
| **Ku::Ku** | Rv0937c | Complement | Heaton et. al. 2014 |
| **RecA-Ku::Ku** | Rv2737c, Rv0937c | Complement of Ku in the double knockout background | Heaton et. al. 2014 |

## METHODS

### Strains

*Mycobacterium tuberculosis* (Mtb) laboratory reference strains H37Rv and Erdman were obtained from the American Type Culture Collection (ATCC). Genotypic confirmation of strain identity was performed using Oxford nanopore (ONP) next generation sequencing (NGS). All strains including CRISPRi and chromosomal deletion mutants were PDIM+ by direct detection on LC-MS. Mutant strains were generously provided by Dr. Michael Glickman (Heaton 2014). Δ*recA*::*zeo*, ΔKu::*hyg*, ΔRecA::RecA (*zeo, kan)*, ΔKu::Ku (*hyg, kan)*, ΔRecAΔKu (*zeo, hyg*), ΔRecAΔKu::Ku (*zeo, hyg, kan*). *Zeo* = Zeomicin, *hyg* = Hygromycin, *kan* = Kanamycin.

### Culture and desiccation assay

Bacterial strains were cultured in standing tissue culture flasks in a biosafety level 3 facility at 37 °C with 5% supplemental CO_2_ in Middlebrook 7H9 media (BD Biosciences) containing 0.05% Tyloxapol (Sigma-Aldrich), 0.2% (w/v) glucose, 0.5 g/L BSA, 0.5% (v/v) glycerol, and 0.085% (w/v) NaCl. When optical density reached OD_600_ 0.8, 1 mL aliquots were transferred to autoclave-sterilized 0.22 µm PVDF Durapore membrane filters (Thomas Scientific) and media removed by vacuum filtration. Filters were transferred to 7H10 agar containing 0.2% (w/v) glucose, 0.5 g/L BSA and 0.085% (w/v) NaCl and incubated at 37 °C with 5% supplemental CO_2_ for 5 days. Filters were then transferred to modified 6-well plates affixed with autoclave-sterilized caps that served as reservoirs for the desiccation experiment. The reservoirs were left empty for desiccation and filled with 200 mM NaCl (unless otherwise specified based on experimental conditions) for wet controls. The plates were incubated at 37 °C in air held at 70% relative humidity (unless otherwise specified) using a Baker Invivo glovebox. Filter-laden desiccated cells were rehydrated by filling the empty reservoirs with either 200 mM NaCl or 7H9 and continuing incubation at 37 °C and 70% RH for 24 hours before processing. CRISPRi strains and complements required 50 µg/mL kanamycin supplemented to broth, agar and saline solution to maintain the plasmid. Cultures were passaged a maximum of 3 times before discarding and returning to frozen stock.

### MAF adaptation and desiccation assay

Model aerosol fluid (MAF) was prepared as previously described (*16*). H37Rv grown to mid-log phase was resuspended in MAF at an OD of 1.0. The cell suspension was incubated in standing culture under hypoxic atmosphere (O_2_ 0.2%, CO_2_ 5%) at 37 °C for two weeks followed by microaerophilic atmosphere (O_2_ 10%, CO_2_ 5%) for an additional two weeks. Upon full reaeration, the cell suspension was arrayed into 2 µL droplets on a polystyrene surface using an Integra Viaflo 384 automated pipette and allowed to desiccate in air at 25 °C in a chamber containing Dreirite to maintain a RH of approximately 0%. Following 24 hours of desiccation, dried residues were rehydrated into 15 mL of MEM-α per plate for 20 minutes after which cells were scraped from the plate, resuspended in Trizol, and processed for RNA Seq as below.

### CFU and limiting dilution assay

Cell-laden filters were plunged into 1 mL phosphate buffered saline (PBS) containing 0.05% tyloxapol (PBST) and 200 µL of 0.1 mm Zirconia silica beads (Fisher Scientific). Gentle bead beating at 2800 rpm for 10 seconds was used to scrape the cells from the filter, de-clump and resuspend. The filter was removed with tweezers and the tubes were allowed to stand still for 10 minutes to allow debris to settle. An aliquot of the resulting cell suspension was serially diluted into PBST and plated on 7H10 agar. Colonies were enumerated after 21 days of incubation at 37 °C in air. A limiting dilution assay was performed following a previously published protocol (*19*).

### RNA Seq

Cell-laden filters were plunged into 2 mL screw cap threaded tubes with O-ring seal (Sarstedt) containing cold 1 mL Trizol (Sigma-Aldrich) and 200 µL of 0.1 mm Zirconia silica beads (Fisher Scientific). Samples were stored at −80 °C until ready for processing. To lyse cells, tubes were bead beat using a Precellys cell homogenizer equipped with air cooling set to 6000 rpm x 30 seconds x 3 cycles with 30 second rest periods in between to prevent sample overheating. After centrifugation lysates were transferred to a new 1.7 mL microfuge tube (Thomas Scientific) and extracted with 250 µL molecular biology grade chloroform (Sigma-Aldrich) and removed from the biosafety level 3 facility for further processing. The aqueous phase was separated, total RNA was precipitated by isopropyl alcohol and washed with ethanol. DNA was depleted with Turbo DNAse (Thermo Fisher) and RNA purified using Zymo clean and concentrator (Zymo Research) according to manufacturer specifications. Prepared libraries were sequenced with paired-end 100 bps on a NovaSeq6000 sequencer. Raw reads were processed using Illlumina bcl2fastq (v2.20) and trimmed with cutadapt (v1.18). Alignment to the H37Rv Mtb reference genome was done using Bowtie2 (v2.2.8) (*56*). Read counts per gene were extracted using HTSeq-count (v0.11.2) and differential expression analysis was generated with DESeq2 (*57, 58*). Pairwise comparisons between groups were generated using parametric tests following a negative binomial distribution with a gene-specific dispersion parameter. Corrected p-values were calculated using the Benjamini-Hochberg multiple hypothesis correction. PCA plots were generated and visualized using tidyverse (v2.0.0) and ggplot2 (v3.5.1) using custom R-scripts.

### Metabolomics

Cell-laden filters were plunged into 2 mL screw cap threaded tubes with O-ring seal (Sarstedt) containing 1mL of a 2:2:1 HPLC-grade Methanol:acetonitrile:water (Sigma-Aldrich) solution. Cells were lysed in an air-chilled Precellys homogenizer at 6000 rpm x 3 cycles x 30 seconds per cycle with 30 seconds rest in between to prevent sample overheating. Lysates were filtered through a 0.22 µm column to ensure sterilization before removal from BSL3 laboratory. 100 µL aliquot of lysate was combined with 100 µL of ddH_2_O containing 0.2% formic acid. The metabolite admixture was chromatographically separated using a Cogent Diamond Hydride type C column using a gradient mobile phase (flow rate 0.4 mL/min) composed of Solvent A (ddH_2_O with 0.2% formic acid) and Solvent B (acetonitrile with 0.2% formic acid). Starting composition of solvents was 15% Solvent A and 85% Solvent B and rising to 95% Solvent A and 5% Solvent B over 24 minutes. Following chromatography, the sample underwent electrospray ionization on an Agilent 6546 ESI-QTOF. Dry gas temperature was set to 350 °C, gas flow 10 L/min, nebulizer pressure 35 psi, and capillary voltage of 3500 V. Ions were detected in both positive mode and negative mode (m/z range: 50 – 1700). Internal calibrants (positive: 64.0472 and 922.009798, negative: 62.032683 and 983.035752) were infused continuously. For charged metabolites such as nucleotide phosphates hydrophilic interaction chromatography (HILIC) was used by separating the extract on a Poroshell 120 Hilic-Z column (150 mm) using a gradient mobile phase (flow rate 0.25 mL/min) composed of Solvent C (10mM ammonium acetate in ddH_2_O with 5 µM Infinity Lab deactivator (Agilent)) and Solvent D (95:15 Acetonitrile:ddH_2_O with 10 mM ammonium acetate and 5 µM Infinity Lab deactivator). Starting composition of solvents was 4% Solvent C and 96% Solvent D and rising to 35% Solvent C and 65% Solvent D over 24 minutes. Following chromatography the sample underwent electrospray ionization on an Agilent Revident ESI-QTOF. Dry gas temperature was set to 300 °C, gas flow 3.5 L/min, nebulizer pressure 35 psi, and capillary voltage of 3500 V. Ions were detected in negative mode (m/z range: 50 – 1700). Internal calibrants 62.032683 and 983.035752 were infused continuously. Metabolites were identified using unique mass-retention time identifiers for masses exhibiting the expected isotopic distribution. An in-house admixture of metabolites spiked into a mycobacterial metabolite matrix was used to confirm chromatographic and analytical integrity. Metabolite abundance was determined by integrating total ion counts over a retention time range using Profinder 6.0 and Qualitative Analysis 6.0 (Agilent Technologies).

### CellRox

Cell-laden filters were plunged into 2 mL screw cap threaded tubes with O-ring seal (Sarstedt) containing 1 mL PBST and 200 µL of 0.1 mm Zirconia silica beads (Fisher Scientific). Gentle bead beating for 10 seconds was used to scrape the cells from the filter and resuspend. Samples were treated with 1 µM CellRox Green Reagent (Thermo Scientific) and incubated at 37 °C with 5 % supplemental CO_2_ for 60 minutes. Cells were washed with PBST x2 times, then resuspended in 4% paraformaldehyde to fix overnight. PFA was removed by washing with PBST. Cells were resuspended in 500 µL of PBST and analyzed with a BD FACSCelesta flow cytometer equipped with BD FACSDiva software. Voltages were adjusted to capture a dynamic range of unstained controls against high signal samples. At least 5,000 events were captured for each sample. Data were analyzed using FCS Express 7.

### LPO

Lipid peroxides were measured using the Lipid hydroperoxide (LPO) quantification kit (Cayman Chemical). Cell-laden filters were plunged into 2 mL screw cap threaded tubes with O-ring seal (Sarstedt) containing 1 mL PBST and 200 µL of Zirconia silica beads (Fisher Scientific). Gentle bead beating for 10 seconds was used to scrape the cells from the filter and resuspend. Cells were resuspended in ddH_2_O and combined with an equal volume of saturated Extract R in molecular biology grade methanol (Sigma Aldrich). Deoxygenated chloroform was prepared by bubbling nitrogen gas through for 10 minutes and then combined with Extract R-treated cells to extract lipids. Lipid extracts were diluted to 950 µL using a 2:1 chloroform:methanol solution and treated with 50 µL of chromogen (1:1 mixture of FTS reagent 1 and reagent 2). A positive control was included in which log-phase cells were treated with 10 mM cumene hydrogen peroxide (Sigma Aldrich) for 1 hr at 37 °C. A negative control was prepared and consisted of CHP-unexposed log phase cell lipid extract without chromogen. Absorbance was read at 500 nm using a SpectraMax M2 spectrophotometer. A lipid peroxide library was used to construct a standard curve. Absorbances were normalized to each samples protein content as determined by Bradford assay.

### Terminal deoxynucleotidyl transferase dUTP nick end labeling (TUNEL) Assay

Single- and double-stranded DNA breaks were detected using the APO-DIRECT kit (BD Biosciences). Cell-laden filters were plunged into 1 mL of 10mM (Dioctyl sulfosuccinate sodium salt) AOT in heptane and agitated to remove cells from the filter. The filter was removed with tweezers and the cell-AOT suspension was placed on ice and inverted every 5 minutes for 1 hour to remove the outer mycobacterial membrane. After pelleting, the cells were fixed with 4% PFA and stored at 4 °C overnight. Fixed cells were washed with PBST and split into two tubes. One set was treated with TdT enzyme, reaction buffer containing bromouracil nucleotides, FITC-labeled antibody and incubated for 1 hour at 37 °C in the dark. The other was treated in kind, but without enzyme to serve as negative controls. After washing with PBST, cells were treated with propidium iodide (PI) at RT for 30 minutes in the dark. One negative TUNEL control was split and used to generate a TUNEL-/PI-sample. Cells were suspended in PI/RNAse solution and analyzed by flow cytometry. Cells staining was quantified using a BD FACSCelesta flow cytometer equipped with BD FACSDiva software. Voltages were adjusted to capture a dynamic range of unstained controls against high signal samples. At least 5,000 events were captured for each sample. Cells were considered positively staining if they exhibited a signal greater than the 99^th^ percentile of unstained controls. Data were analyzed using FCS Express 7.

### CRISPRi strain construction and validation

Small guide RNA oligomers were designed using the Pebble database managed by Rockefeller University (pebble.rockefeller.edu) and purchased from Integrated DNA technologies (IDT). Parent plasmid PLJR965 was digested with the restriction enzyme BsmBI (NEB) and sgRNA oligomers ligated with T4 ligase (NEB) in a one-pot synthesis. Mtb gene-targeting plasmids were transformed into DH5a competent *E. col*i using heat shock and transformants were selected on kanamycin selection agar. Plasmid DNA from colony-derived culture was isolated for transformation into Mtb using a Qiagen miniprep kit Electroporation into competent H37Rv and plating on kanamycin selective 7H10 agar yielded Mtb strains capable of ATC-dependent targeted gene silencing. Appropriate sequence positioning and alignment was confirmed by oxford nanopore NGS.

### RT qPCR for fold-change

Cell-laden filters were plunged into 1.7 mL threaded tubes with O-ring seal containing cold 1 mL Trizol (Aldrich chemical) and 200 µL of 0.1 mm Zirconia silica beads (Fisher Scientific). Samples were stored at −80 °C until ready for processing. To lyse cells, tubes were bead beat using a cell homogenizer equipped with air cooling set to 6000 rpm x 30 seconds x 3 cycles with 30 second rest periods in between to prevent sample overheating. Lysates were passed through a 0.2 µm filter to ensure sterilization, extracted with 250 µL molecular biology grade chloroform and removed from the biosafety level 3 facility for further processing. RNA was isolated using Directzol RNA MiniPrep (Zymo research). DNA in the eluant was depleted by incubating with 4U DNAse (Invitrogen) at 25 °C for 1 hour. The reaction was quenched with DNAse inactivator (Invitrogen) and purified once again using Directzol RNA MiniPrep. After confirming purity by nanodrop, 10 µL of 100 ng/mL RNA was converted to cDNA by incubating with 4U/µL Maxima H Reverse transcriptase (ThermoFisher) for 10 minutes at 25 °C, 30 minutes at 60 °C followed by heat inactivation for 5 minutes at 85 °C. Amplification of products was performed using forward and reverse primers (IDT) and Taq polymerase Sybr green master mix (ThermoFisher) on a Lightcycler 480 II (Roche). Each PCR cycle included 95 °C for 30s, 60 °C for 30s, and 72 °C for 20s for a total of 45 cycles. Crossing points were determined using Lightcycler 480 SW software (v1.5.1). Amplification of the housekeeping gene *sigA* was used to calculate the ΔΔCp and fold change was calculated according to the following equation: FC = 2^(-ΔΔCp). For desiccation-rehydration survival screens, the following equation was used to calculate relative survival:

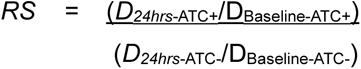

*RS* = Relative Survival

D_24hrs_ = CFU/mL after 24 hours drying

D_Baseline_ = CFU/mL at Baseline

### Mutation frequency analysis

Mutation frequency was calculated using a Luria-Delbrück fluctuation assay. Cell-laden filters were processed the same as for CFU except they were additionally plated on 7H10 agar containing 1 µg/mL rifampin or 1.5 µg/mL ciprofloxacin. Colonies were enumerated after 4 weeks incubation at 37 °C in air. FALCOR, the online tool for fluctuation analysis (genomics.brocku.ca/FALCOR) was used to determine mutation frequency using the Lea-Coulson method.

### rpoB PCR and SNP analysis

Rif-R colonies were picked at random and expanded in 7H9 media to a volume of 5 mL. At OD_600_ = 1.0 the cells were pelleted and resuspended in 1mL PBST then transferred to O-ring tubes containing 200 µL of 0.1 mm Zirconia silica beads (Fisher Scientific). Cells were lysed using an air-chilled Percellys homogenizer at 6000 rpm x 30s x 3 cycles with 30s cool down in between cycles to prevent overheating. Filter sterilization through spin-X columns allowed removal from BSL3. gDNA was isolated using a phenol-chloroform-isoamyl alcohol extraction, followed by isopropanol and sodium acetate precipitation. Samples were resuspended in ddH_2_O and purity was confirmed by Nanodrop. Amplification of 1 µL gDNA using *rpoB* primers and 2U/mL Q5 DNA polymerase with Q5 High-GC followed by QiaQuick nucleotide cleanup Kit yielded 50 – 200 ng/µL of cDNA. Sequencing of the amplicons was completed using ONP NGS and alignments to WT *rpoB* were made in SnapGene.

### Transmission simulation system (TSS)

CRISPRi strains targeting *mfd* with either Rif-S WT *rpoB* or Rif-R S450L *rpoB* were cultured in 7H9 containing 50 µg/mL kanamycin +/- 500 ng/µL ATC for 10 generations. Cells were kept in mid-log phase by serial passaging. ATC was replenished every 3 – 4 days and at passage. Cells were suspended in 5 mL PBS and nebulized into the TSS via a single-jet Blaustein atomizer module (CH technologies) using a controlled flow rate of 12 mL/hr delivered via syringe pump. A scattered-light aerosol spectrometer (Promo 2000 with Aerosol Sensor welas 2070, Palas) for particle concentration monitoring and a 6-stage viable Andersen cascade impactor (Tisch Environmental) were equipped to monitor particle size distribution. Particles were collected into an impinger containing PBS and plated onto Middlebrook 7H10 agar to enumerate biomass by CFU.

### Clinical strain phylogenies

Genomic analysis of the 51,229 strains of Mtb clinical isolates circulating globally and the 494 isolates from India have been used in previous reports (*41, 42*). Briefly, phylogenetic reconstruction was achieved by combining all SNP locations for each isolate into a consensus list, aligned using MEGA 6.0 and visualized in FigTree (version 1.4.3). Each dataset was used to identify strains with non-synonymous SNPs within *mfd* and known Rif-R conferring SNPs within *rpoB* to calculate ratios of specific alleles within subpopulations and clades.

## Notes

### Competing Interest Statement

The authors have declared no competing interest.

## References

1. G. A. Goig et al., Ecology, global diversity and evolutionary mechanisms in the Mycobacterium tuberculosis complex. Nat Rev Microbiol, (2025).

2. G. A. Goig et al., Transmission as a Key Driver of Resistance to the New Tuberculosis Drugs. N Engl J Med 392, 97–99 (2025).

3. Global tuberculosis report 2024. Geneva: World Health Organization. (2024).

4. J. D. Ernst, The immunological life cycle of tuberculosis. Nat Rev Immunol 12, 581–591 (2012).

5. C. J. Cambier, S. Falkow, L. Ramakrishnan, Host evasion and exploitation schemes of Mycobacterium tuberculosis. Cell 159, 1497–1509 (2014).

6. P. Chandra, S. J. Grigsby, J. A. Philips, Immune evasion and provocation by Mycobacterium tuberculosis. Nat Rev Microbiol 20, 750–766 (2022).

7. R. L. Riley et al., Aerial dissemination of pulmonary tuberculosis. A two-year study of contagion in a tuberculosis ward. 1959. Am J Epidemiol 142, 3–14 (1959).

8. K. P. Fennelly et al., Variability of infectious aerosols produced during coughing by patients with pulmonary tuberculosis. Am J Respir Crit Care Med 186, 450–457 (2012).

9. B. Patterson et al., Cough-independent production of viable Mycobacterium tuberculosis in bioaerosol. Tuberculosis (Edinb) 126, 102038 (2021).

10. R. Dinkele et al., Capture and visualization of live Mycobacterium tuberculosis bacilli from tuberculosis patient bioaerosols. PLoS Pathog 17, e1009262 (2021).

11. V. Nduba et al., Mycobacterium tuberculosis cough aerosol culture status associates with host characteristics and inflammatory profiles. Nat Commun 15, 7604 (2024).

12. S. J. Webb, FACTORS AFFECTING THE VIABILITY OF AIR-BORNE BACTERIA: I. BACTERIA AEROSOLIZED FROM DISTILLED WATER. Canadian Journal of Microbiology 5, 649–669 (1959).

13. S. J. Webb, Factors affecting the viability of air-borne bacteria. III. The role of bonded water and protein structure in the death of air-borne cells. Can J Microbiol 6, 89–105 (1960).

14. R. G. Loudon, L. R. Bumgarner, J. Lacy, G. K. Coffman, Aerial transmission of mycobacteria. Am Rev Respir Dis 100, 165–171 (1969).

15. M. Potts, Desiccation tolerance of prokaryotes. Microbiological Reviews 58, 755–805 (1994).

16. S. Mishra et al., Candidate transmission survival genome of Mycobacterium tuberculosis. Proc Natl Acad Sci U S A 122, e2425981122 (2025).

17. H. Eoh et al., Metabolic anticipation in Mycobacterium tuberculosis. Nat Microbiol 2, 17084 (2017).

18. R. S. Jansen, K. Y. Rhee, Emerging Approaches to Tuberculosis Drug Development: At Home in the Metabolome. Trends Pharmacol Sci 38, 393–405 (2017).

19. K. Saito et al., Rifamycin action on RNA polymerase in antibiotic-tolerant Mycobacterium tuberculosis results in differentially detectable populations. Proc Natl Acad Sci U S A 114, E4832–e4840 (2017).

20. K. Saito et al., Oxidative damage and delayed replication allow viable Mycobacterium tuberculosis to go undetected. Sci Transl Med 13, eabg2612 (2021).

21. S. K. Hatzios et al., Osmosensory signaling in Mycobacterium tuberculosis mediated by a eukaryotic-like Ser/Thr protein kinase. Proc Natl Acad Sci U S A 110, E5069–5077 (2013).

22. N. Robinson et al., Mycobacterial phenolic glycolipid inhibits phagosome maturation and subverts the pro-inflammatory cytokine response. Traffic 9, 1936–1947 (2008).

23. M. V. Claudia, A. A. Javiera, N. S. Sebastián, F. R. José, L. Gloria, Interplay between desiccation and oxidative stress responses in iron-oxidizing acidophilic bacteria. J Biotechnol 383, 64–72 (2024).

24. H. P. Oswin et al., Oxidative Stress Contributes to Bacterial Airborne Loss of Viability. Microbiol Spectr 11, e0334722 (2023).

25. H. P. Oswin et al., An assessment of the airborne longevity of group A Streptococcus. Microbiology (Reading) 170, (2024).

26. L. Contreras-Porcia, D. Thomas, V. Flores, J. A. Correa, Tolerance to oxidative stress induced by desiccation in Porphyra columbina (Bangiales, Rhodophyta). J Exp Bot 62, 1815–1829 (2011).

27. M. I. Goncheva, D. Chin, D. E. Heinrichs, Nucleotide biosynthesis: the base of bacterial pathogenesis. Trends Microbiol 30, 793–804 (2022).

28. J. M. Rock et al., Programmable transcriptional repression in mycobacteria using an orthogonal CRISPR interference platform. Nat Microbiol 2, 16274 (2017).

29. R. Gupta, D. Barkan, G. Redelman-Sidi, S. Shuman, M. S. Glickman, Mycobacteria exploit three genetically distinct DNA double-strand break repair pathways. Mol Microbiol 79, 316–330 (2011).

30. E. M. Witkin, Radiation-induced mutations and their repair. Science 152, 1345–1353 (1966).

31. E. M. Witkin, Mutation frequency decline revisited. Bioessays 16, 437–444 (1994).

32. C. P. Selby, E. M. Witkin, A. Sancar, Escherichia coli mfd mutant deficient in “mutation frequency decline” lacks strand-specific repair: in vitro complementation with purified coupling factor. Proc Natl Acad Sci U S A 88, 11574–11578 (1991).

33. C. P. Selby, A. Sancar, Molecular mechanism of transcription-repair coupling. Science 260, 53–58 (1993).

34. O. Adebali et al., The Mfd protein is the transcription-repair coupling factor (TRCF) in Mycobacterium smegmatis. J Biol Chem 299, 103009 (2023).

35. B. Martinez, B. K. Bharati, V. Epshtein, E. Nudler, Pervasive Transcription-coupled DNA repair in E. coli. Nat Commun 13, 1702 (2022).

36. E. Nudler, Transcription-coupled global genomic repair in E. coli. Trends Biochem Sci 48, 873–882 (2023).

37. M. N. Ragheb et al., Inhibiting the Evolution of Antibiotic Resistance. Mol Cell 73, 157–165.e155 (2019).

38. <WHO 2021 report of known mutations.pdf>.

39. S. Gagneux et al., The competitive cost of antibiotic resistance in Mycobacterium tuberculosis. Science 312, 1944–1946 (2006).

40. K. A. Eckartt et al., Compensatory evolution in NusG improves fitness of drug-resistant M. tuberculosis. Nature 628, 186–194 (2024).

41. Q. Liu et al., Tuberculosis treatment failure associated with evolution of antibiotic resilience. Science 378, 1111–1118 (2022).

42. S. K. Shanmugam et al., Mycobacterium tuberculosis Lineages Associated with Mutations and Drug Resistance in Isolates from India. Microbiol Spectr 10, e0159421 (2022).

43. J. Y. Kang et al., Structural basis for transcription complex disruption by the Mfd translocase. Elife 10, (2021).

44. E. Mansour et al., Measurement of temperature and relative humidity in exhaled breath. Sensors and Actuators B: Chemical 304, 127371 (2020).

45. H. C. Leyva-Sánchez et al., Role of Mfd and GreA in Bacillus subtilis Base Excision Repair-Dependent Stationary-Phase Mutagenesis. J Bacteriol 202, (2020).

46. J. Carvajal-Garcia, A. N. Samadpour, A. J. Hernandez Viera, H. Merrikh, Oxidative stress drives mutagenesis through transcription-coupled repair in bacteria. Proc Natl Acad Sci U S A 120, e2300761120 (2023).

47. B. D. Kana et al., Role of the DinB homologs Rv1537 and Rv3056 in Mycobacterium tuberculosis. J Bacteriol 192, 2220–2227 (2010).

48. P. Dupuy, M. Howlader, M. S. Glickman, A multilayered repair system protects the mycobacterial chromosome from endogenous and antibiotic-induced oxidative damage. Proc Natl Acad Sci U S A 117, 19517–19527 (2020).

49. P. Dupuy et al., Distinctive roles of translesion polymerases DinB1 and DnaE2 in diversification of the mycobacterial genome through substitution and frameshift mutagenesis. Nat Commun 13, 4493 (2022).

50. P. Dupuy et al., Roles for mycobacterial DinB2 in frameshift and substitution mutagenesis. Elife 12, (2023).

51. B. E. Heaton, D. Barkan, P. Bongiorno, P. C. Karakousis, M. S. Glickman, Deficiency of double-strand DNA break repair does not impair Mycobacterium tuberculosis virulence in multiple animal models of infection. Infect Immun 82, 3177–3185 (2014).

52. C. Loiseau et al., The relative transmission fitness of multidrug-resistant Mycobacterium tuberculosis in a drug resistance hotspot. Nat Commun 14, 1988 (2023).

53. M. Silcocks et al., Evolution and transmission of antibiotic resistance is driven by Beijing lineage Mycobacterium tuberculosis in Vietnam. Microbiol Spectr 11, e0256223 (2023).

54. Q. Liu et al., Local adaptation of Mycobacterium tuberculosis on the Tibetan Plateau. Proc Natl Acad Sci U S A 118, (2021).

55. T. Hulsen, DeepVenn -- a web application for the creation of area-proportional Venn diagrams using the deep learning framework Tensorflow.js. arXiv, (2022).

56. B. Langmead, S. L. Salzberg, Fast gapped-read alignment with Bowtie 2. Nat Methods 9, 357–359 (2012).

57. S. Anders, P. T. Pyl, W. Huber, HTSeq--a Python framework to work with high-throughput sequencing data. Bioinformatics 31, 166–169 (2015).

58. M. I. Love, W. Huber, S. Anders, Moderated estimation of fold change and dispersion for RNA-seq data with DESeq2. Genome Biol 15, 550 (2014).

